# ARF degradation defines a deeply conserved step in auxin response

**DOI:** 10.1101/2024.09.12.612472

**Authors:** Martijn de Roij, Jorge Hernández García, Shubhajit Das, Jan Willem Borst, Dolf Weijers

**Affiliations:** Laboratory of Biochemistry, Wageningen University, Stippeneng 4, 6708WE Wageningen, the Netherlands

**Author notes:** Institute for Science and Technology Austria, Klosterneuburg, Austria (Shubhajit Das).

## Abstract

Auxin response critically depends on the concentrations and stoichiometry of competing A- and B-class AUXIN RESPONSE FACTOR (ARF) proteins. In *Marchantia polymorpha*, both A- and B-ARFs are unstable, and here we identify a minimal necessary and sufficient region for ARF degradation that is critical for development, and auxin response. Through comparative analysis, we find that ARF instability likely preceded the emergence of the auxin response system.

## Main text

Auxin is a central signalling molecule in plant development^1^ and acts chiefly through the activation of Auxin Response Factors (ARFs), a family of DNA-binding transcription factors^2^. ARFs are phylogenetically divided into three classes (A, B, and C)^3^. A-ARFs are auxin-dependent gene regulators and are antagonized by auxin-independent B-ARFs^4,5^. Based on work in the liverwort *Marchantia polymorpha*, C-ARFs appear detached from auxin response^4^. Given that competition between A- and B-ARFs determines auxin response, their levels and stoichiometry are key parameters defining this response. We recently developed fluorescent knock-in reporters of all auxin response proteins in Marchantia^6^, which encodes single copies of each ARF class. We found that both A- (MpARF1) and B-class (MpARF2) ARFs are unstable, and that this instability requires the 26S proteasome^6^. Several other ARFs have been reported to be target for targeted degradation (reviewed in Ref. 7), but many questions regarding mechanisms, biological significance and evolutionary origin of the instability remain.

To address these questions, we first mapped the region in MpARF2 conferring instability. We expressed mNeonGreen (mNG) fluorescently tagged, nuclear targeted, protein domains from the native *MpARF2* promoter (Fig. 1a). While the NLS-mNG control accumulated to high levels, full-length MpARF2 (NLS-FL-mNG) could not be detected (Fig. 1a,b,d) unless the 26S proteasome was inhibited with Bortezomib (Bz; Fig. 1c), as expected. While mNG fusions to the Middle Region (NLS-MR-mNG) or PB1 domains (NLS-PB1-mNG) were stable (Fig. 1a,b,d), the DNA-Binding Domain (NLS-DBD-mNG) alone was sufficient to confer instability (Fig. 1a,b,d). Again, Bz treatment led to protein accumulation (Fig. 1c,d). Thus, the MpARF2 DBD contains the minimal elements required for instability.

**Figure 1:**
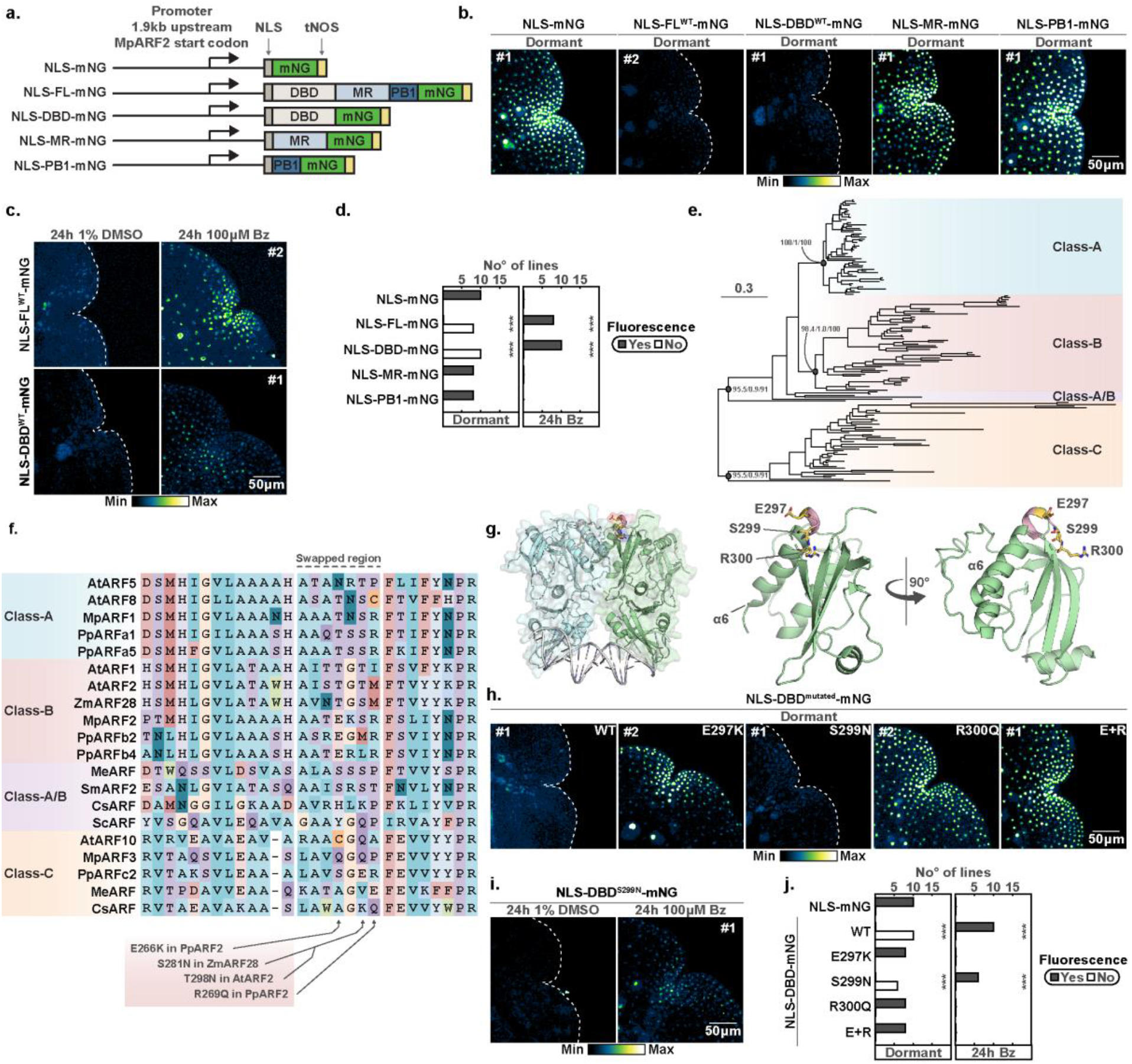
Identification of a motif in MpARF2 essential for instability. **a**. Schematic overview of translational fusion proteins assessed in subsequent assays. **b**. Confocal images of *M. polymorpha* gemmae, showing expression of fusion proteins outlined in a. **c**. Gemmae were treated with Bz or DMSO and imaged by confocal microscopy, proteins correspond to a. **d**. Screening of nuclear fluorescence of independent transgenic lines, if no fluorescence in dormant gemmea was detected (left panel), plants were treated with Bz and fluorescence was reassessed (right panel). Fluorescence occurance was compared to NLS-mNG via a chi-squared statistical test (*** *p*<0.001). **e**. Phylogenetic relationships between ARFs. Statistical support for major nodes are shown as SH-aLRT branch test/Approximate bayesian test/Ultrafast bootstrap approximation values. **f**. Multiple sequence alignment of ARFs representing the major classes. Putative amino acids responsible for instability (and the swapped regions) are outlined. **g**. MpARF2 DBD crystal structure with key residues E297, S299, and R300 highlighted and enlarged. **h**. Confocal images of gemmae expressing NLS-DBD-mNG fusions carying outlined mutations. **i**. Same as c. **j**. Same as d., similar controls were plotted.. **c.-d. & i.-j**. Bz, Bortezomib. **b.-c. & h.-i**. Independent transgenic lines denoted by numbers with a hash, one representative line is shown.

Recently, mutations in the DBD of *Physcomitrium patens* and *Zea mays* (Maize) B-ARFs were shown to inhibit proteasome-dependent degradation. In both cases, mutations map to the same short motif and engineering an equivalent mutation in *Arabidopsis thaliana* AtARF2 also reduced breakdown^8^. It is therefore possible that instability in B-ARFs has a single, common origin in ARF evolution (Fig. 1e). The existence of a single B-ARF in Marchantia enables testing this hypothesis. Indeed, the amino acids required for instability in Physcomitrimum and Maize are conserved in MpARF2 (E297, S299, and R300 in MpARF2; Fig. 1f). When mapped onto our experimentally determined MpARF2 DBD structure, we found these to be positioned in an outward facing loop towards the C-terminus of α-helix 6, likely mediating DBD homodimerization (Fig. 1f-g). We engineered E297K, S299N, and R300Q single mutations, as well as E297K+R300Q (E+R) double mutations in the NLS-DBD-mNG protein and analyzed accumulation and stability. All mutations except S299N led to strong nuclear signal (Fig. 1h-j; SI Fig. 1f-g). In contrast, the S299N mutation did not stabilize the protein (Fig. 1h-j; SI Fig. 1f-g). Thus, B-ARF degradation likely has a single evolutionary origin, but as discussed in^8^, there probably is co-evolution between the ARF and its proteolysis partner.

We deliberately engineered mutations in the DBD, to uncouple degradation and accumulation from potential effects on plant growth and development. It is however unclear what biological relevance MpARF2 degradation has. To address the impact of a lack of MpARF2 degradation on growth, development and auxin response, we engineered the E+R mutation in the full-length fusion protein. As expected, NLS-FL^E+R^-mNG-bearing gemmae exhibited very high levels of nuclear mNG fluorescence whereas lines expressing NLS-FL^WT^-mNG did not unless they were treated with Bz (Fig. 2a; Fig. 1b-c; SI Fig. 1d). MpARF2-accumulating lines displayed a wide range of strong developmental defects, none of which were observed in NLS-FL^WT^-mNG lines (SI Fig. 4). Most NLS-FL^E+R^-mNG lines were dwarfed, showed epinastic growth, ectopic apical notch formation, and failed to produce gemmae (SI Fig. 4). Some lines did form gemmae (#7 and #10), even though they mostly did so on the thallus, rather than inside gemmae cups (SI Fig. 5b). However, these lines allowed us to explore effects of MpARF2 accumulation on gemmae development and auxin response. Mature gemmae showed defects in apical (meristematic) notch formation, often showing many more notches than the regular two notches (Fig. 2a; SI Fig. 4c). EdU staining of S-phase cells confirmed the existence of supernumerary notches (Fig. 2b). As expected from B-ARF accumulation, NLS-FL^E+R^-mNG gemmae showed a strong reduction in auxin response (Fig. 2c-d; SI Fig. 5d). Thus, regulation of MpARF2 levels through degron-mediated proteasomal degradation is necessary for normal development and auxin response.

**Figure 2:**
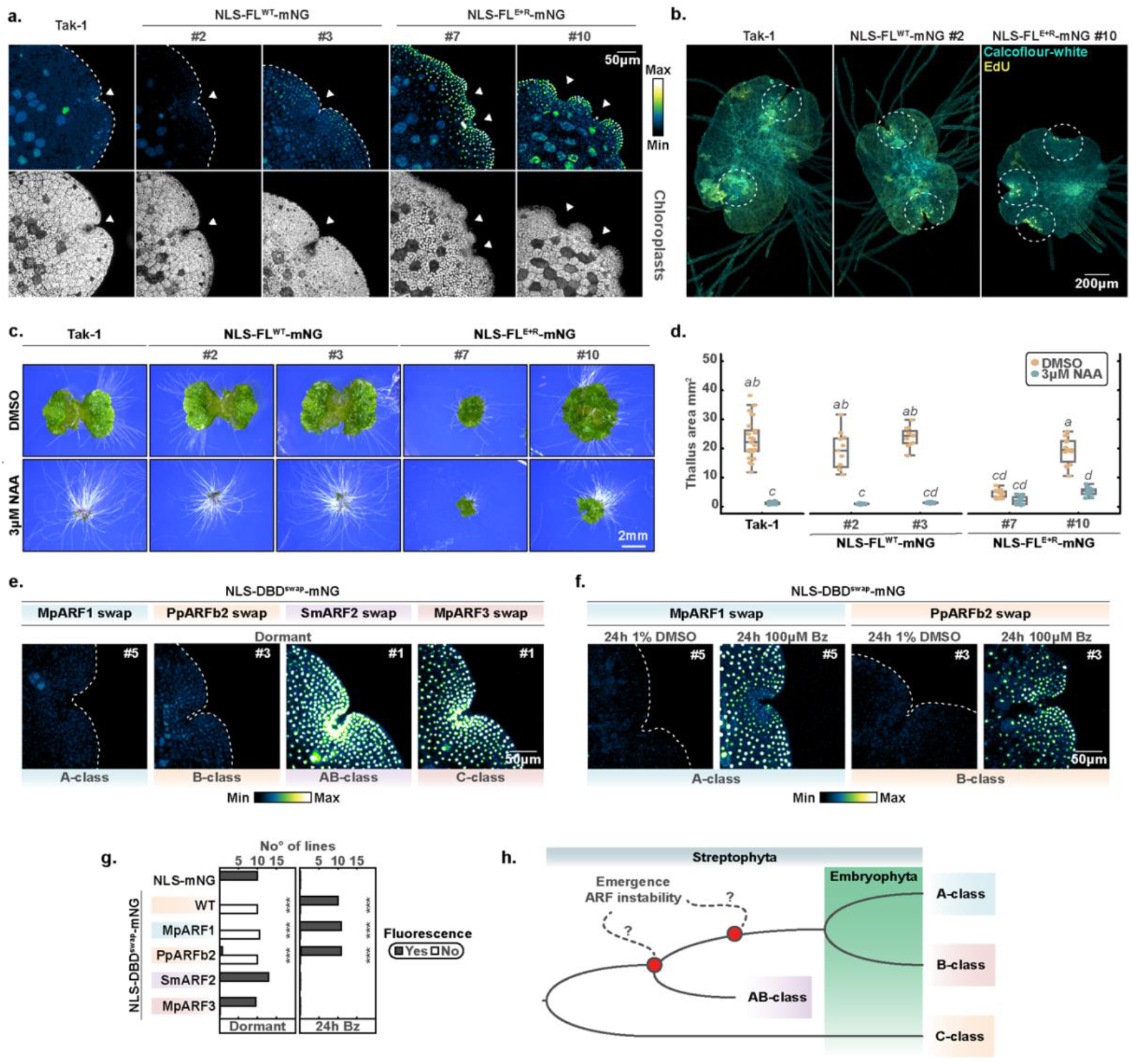
Biological significance of MpARF2 degradation and the evolutionary origin of ARF instability. **a**. Confocal images of Tak-1 wildtype -and transgenic gemmae expressing NLS-FL^WT^-mNG or NLS-FL^E+R^-mNG constructs. Chloroplast autofluorescence outlines gemmae morphology. Hashed numbers are independent transgenic lines. **b**. Confocal images of EdU staining in 3-day old gemmalings of S-phase nuclei (yellow), cell walls were stained with calcofluor-white (turquoise). **c**. Qualitative comparison of auxin response in gemmalings grown for 10 days on medium containing 3 μM NAA or DMSO, respectively. **d**. Quantification of projected thallus area of gemmalings shown in c. compared per one-way ANOVA with Tukey post-hoc test (italics denote significant differences, *p*<0.05). **e**. Fluorescence accumulation (assessed by confocal microscopy) of swaps of homologous regions of different ARFs (representing major ARF classes) into NLS-DBD-mNG. **f**. Lines which showed no detectable fluoresence in dormant gemmae in e. were treated for 24h with Bz or DMSO and fluorescence accumulation was reassessed. **g**. Screening of nuclear fluorescence of independent transgenic lines, if no fluorescence in dormant gemmae was detected (left panel), plants were treated with Bz and fluorescence was reassessed (right panel). Fluorescence occurrence was compared to NLS-mNG via a chi-squared statistical test (*** *p*<0.001). **h**. Schematic cladogram illustrating the hypothesized evolutionary relationship between known ARF clades and the emergence of the instability-conferring region.

Hence, equivalent mutations stabilize Marchantia, Physcomitrium, Maize, and Arabidopsis ARFs. We next tested if these regions are functionally equivalent by swapping this region from Physcomitrium PpARFb2 into the MpARF2 DBD. The NLS-DBD^PpARF2 swap^-mNG protein was unstable, yet accumulated when treated with Bz (Fig. 2e-g), and the degradation mechanism is thus likely homologous between species.

Given that in Marchantia, both A- and B-ARFs are unstable^6^, and having identified the minimal region that is both necessary and sufficient for MpARF2 degradation, we next asked if mechanisms of instability are universal among A- and B-ARFs. There is some conservation between MpARF1 and MpARF2 in the region required for MpARF2 (and Physcomitrium, Maize and Arabidopsis) instability (Fig. 1f). We tested if the equivalent region from MpARF1 could render MpARF2 unstable, i.e. we swapped this short region into the MpARF2 NLS-DBD-mNG (Fig. 1f). As a control, we included the equivalent region from the C-class MpARF3 which is stable in its native context^6^. While the NLS-DBD^MpARF3 swap^-mNG protein accumulated to high levels even in the absence of Bz, we could hardly detect NLS-DBD^MpARF1 swap^-mNG protein (Fig. 2e-g; SI Fig. 6). When treated with Bz however, this protein accumulated to high levels (Fig. 2e-g; SI Fig. 6). Thus, homologous regions in MpARF1 and MpARF2 confer comparable protein instability.

A- and B-ARFs derive from a duplication in an ancestral A/B-ARF gene^3^, and neofunctionalized following this duplication^9^. The finding that the same region in MpARF1 and MpARF2 can confer instability suggests that this feature has a single origin that predates the divergence of A- and B-ARFs. While the nature of the ancestral A/B ARF is not clear, extant representatives of these proteins are found in the sister lineages to land plants, the streptophyte algae^3^. We selected SmARF2 from the alga *Spirogloea muscicola* as representative of the Zygnematophyceae, the lineage most closely related to land plants^10^, and swapped the homologous region from its A/B-ARF (Fig. 1f) into NLS-DBD-mNG. NLS-DBD^SmARF2 swap^-mNG fusions were highly stable, mimicking behaviour of fusion proteins with mutated degron residues (Fig. 2e-g; SI Fig. 6). This result is consistent with two interpretations: either the ancestral state of the A/B-ARF included protein instability conferred by this region, followed by subsequent loss in the lineage giving rise to SmARF2. Alternatively, the ancestral A/B-ARF state was stable, and instability evolved in the lineage giving rise to land plant prior to A/B divergence (Fig. 1h). The very long evolutionary divergence times in this group of algae (>600 Mya^11^), and sparse species sampling makes such inferences problematic. Further exploration of the equivalent region in a larger number of Streptophyte algal species may help resolve the exact origin of ARF degradation.

Our work identifies a minimal region for ARF degradation in Marchantia and shows that a key element required for instability is conserved among B-ARFs, even in Marchantia which has only a single copy. We show that MpARF2 instability is critical for normal development and auxin response. Our analysis of MpARF1 reveals that the capacity to mediate instability is very likely an ancestral property of the protein that predated the split between A- and B-ARFs. Given that auxin-dependence only evolved after the A/B split^9^, we infer that ARF instability preceded the origin of the auxin response system. A key future question is by what mechanism the minimal region confers instability. This region likely constitutes a protein-protein interaction interface that allows a Ubiquitin ligase, or another proteolysis adaptor protein, to bind. The fact that the S299N mutation does not stabilize MpARF2, while the orthologous mutation stabilizes Maize ARF28, suggests divergence in the degradation interface, and likely co-evolution with a partner protein. We expect that the future identification of other components in ARF degradation will help understanding the mechanisms, diversity, and biological significance of degradation across the many processes that auxin controls.

## Methods

### Sequence identification, alignment and phylogenetic analysis

Most ARF sequences have been previously identified ^3^. Additional sequences have been retrieved through BLASTP analysis using reference databases for *Zea mays* (B73-NAM-5.0.55), *Ceratopteris richardii* (V2.1), *Selaginella kraussiana, Ceratodon purpureus* (GG1), and algal genomes of *Mesotaenium endlicherianum* (SAG12.97), *Chlorokybus melkonianii* (CCAC 0220, previously *C. atmophyticus*), *Zygnema circumcarinatum*, all accessed through Phycocosm. For *M. endlicherianum*, the v2.0 annotation was surveyed to obtain the MeARFab sequence. BLASTP cut-off was set to 10E-10 for algal searches. All sequences were first checked for the presence of ARF DNA binding domain features (*i*.*e*.: B3 DNA binding domain, PF02362; and Auxin_resp, PF06507) or discarded otherwise. Protein sequences were aligned with the *E-INS-i* algorithm of MAFFT version 7 (https://mafft.cbrc.jp/alignment/software/). Alignments were trimmed to keep the DBD sequences and manually curated. TrimAI was used on curated alignments to discard positions with more than 80% gaps (http://trimal.cgenomics.org/). The ModelFinder implementation in IQ-TREE was used on final alignments to choose JTT+I+G4 as substitution model based on the Akaike-and Bayesian Information Criterion. IQ-TREE was also used on the final alignment to build a maximum likelihood phylogenetic tree. Branch statistic support was inferred using Ultrafast bootstrap (5000 alignments), the SH-like approximate Likelihood Ratio Test, and Approximate Bayes Test.

### Structural analysis degron

The MpARF2 crystal structure was retrieved from PDB (https://www.rcsb.org/structure/6SDG)^4^. Structural visualisation was performed using Pymol (v2.3.4) software.

### Plasmid construction

A genomic fragment of 1920 bp upstream of the *MpARF2* startcodon was amplified with primers MdR298 and MdR299 (Table 1). This region was used as promoter to drive expression of the transgenes. Next, the plasmid pMpGWB100, carrying a hygromycin resistance casette, was digested with the restriction enzyme XbaI (ThermoFisher Scientific) and the aforementioned *MpARF2* promoter was ligated downstream of an XbaI site followed by the mNeonGreen (mNG) coding sequence using NEBuilder HiFi DNA Assembly (New England Biolabs, Inc)^12^. Next, MpARF2 domains, the full length protein coding sequence, and mutant versions thereof were amplified from in-house plasmids using primer pairs specified in Table 1 and introduced into the XbaI site flanked by the *MpARF2* promoter and mNG. The degron swap constructs were amplified in two fragments from the MpARF2 DBD with two primer pairs containing a non-complementary sequence to introduce the swap, these fragments were then integrated using HiFi DNA Assembly (SI Table 1). Swaps were generated based on the alignment shown in Fig. 1f.

### Plant growth conditions and transformation

For this study *Marchantia polymorpha* Takaragaike-1 (Tak-1) was used as wild-type. Plants were grown on ½ strength Gamborg B5 medium at 22°C with 40 μmol photons m^−2^s^−1^ continuous white fluorescent light. Tak-1 was transformed using agrobacterium-mediated delivery of transgenic constructs as described by Kubota et al., 2013 ^13^. Transgenic plants were selected on ½ Gamborg B5 with 10 mg/L hygromycin and 100 μg/ml cefotaxime.

### EdU-labeling of S-phase cells

To visualize S-phase cells in 3-day old gemmalings, plants were cultured for 3 h in liquid ½ Gamborg B5 medium containing 20 μM 5-ethynyl-2’deoxyuridine (EdU) followed by fixation in 3.7% formaldehyde in phosphate buffer saline (PBS, pH 7.4) for 1h in a vacuum. Plants were then washed twice in PBS and permeabilized for 20 mins in 0.5% Triton X-100 in PBS. Next, plants were washed twice in PBS containing 3% BSA and placed in the EdU click-IT reaction mixture with the 594 Alexa FLUOR fluorophore in the dark for 1 h (Invitrogen). This was followed by two washes with PBS with 3% BSA after which samples were placed in ClearSee solution for 1-4 days, protected from light^14^. Cell walls were stained using calcofluor-white.

### Confocal microscopy & Assessment of fusion protein stability

All microscopy was performed on a Leica SP8X-SMD confocal microscope fitted with hybrid detectors and a 40 MHz pulsed white-light laser. For mNeonGreen, the fluorophore was excited using 488 nm laser line at 12% power output. Fluorescence was detected between 500-570 nm with hybrid detectors set to photon counting mode with 1.00-24.50 ns time-gating active to suppress background fluorescence. For the EdU staining experiment, 594 Alexa FLUOR was excited with a 594 nm laser line at 9% laser power and fluorescence emission was captured between 600-660 nm (0.7-24.50 ns time-gating active). Z-stacks were acquired using an HC PL APO 20x/0.75 water immersion or APO CS 10x/0.40 dry objective. To display images, ImageJ (v1.52) was used to generate maximum-intensity projections.

When investigating stability of various fusion proteins, we screened T1 transgenic lines for fluorescence by imaging the apical notch region of dormant gemmae. When no fluorescence was detected, we treated these gemmae with Bortezomib (Cayman Chemical) for 24 h or the solvent DMSO (Sigma-Aldrich) as control, and checked for fluorescence again. Transgenic lines that showed no fluorescence before and after Bortezomib treatment were excluded from the analysis.

### Phenotypic analysis

For experiments assessing fluorescence accumulation, non-chimeric transgenic lines were established by growing a G_1_ generation from gemmae of independent plants that appeared resistant to initial selection on hygromycin (T_1_ generation). Gemmae from the G_1_ generation were used for experiments. To assess the phenotypes of plants expressing the full length MpARF2-mNG fusion, we reported on phenotypes of 50 day old T_1_ generation plants. For growth assays, dormant gemmae were micro dissected from the gemmae cups/thallus and if clearly distinguishable notches could be observed, these were counted. Next, the gemmae were placed onto ½ Gamborg B5 medium (in the case of NAA sensitivity, medium was supplemented with DMSO or 3 μM NAA) and grown at 22°C with 40 μmol photons m^−2^ s^−1^ continuous white fluorescent light. After seven (general growth) or ten (NAA sensitivity) days, pictures were taken with a Canon EOS250D camera and thallus area was measured using ImageJ (v1.52). For higher magnification images a Leica M205 stereomicroscope was used. Statistical tests were performed in R (v4.2.1).

## Acknowledgements

We thank Sjoerd Woudenberg for help and advice, André Kuhn for comments on the manuscript, and Michael Prigge and Mark Estelle for helpful discussions

This work was supported by a grant from Netherlands Organization for Scientific Research (NWO; OCENW.M20.031 to J.W.B.), a Marie Skłodowska-Curie Individual Fellowship (H2020-MSCA-IF-2020 to J.H-G.), and a Research Grant from the Human Frontiers Research Program (HFSP; grant RGP0015/2022 to D.W.).

## Competing interests

No competing interests are declared.

## Notes

### Competing Interest Statement

The authors have declared no competing interest.

