## Supplementary File for "ARF degradation defines a deeply conserved step in auxin response"

### Supplementary Information

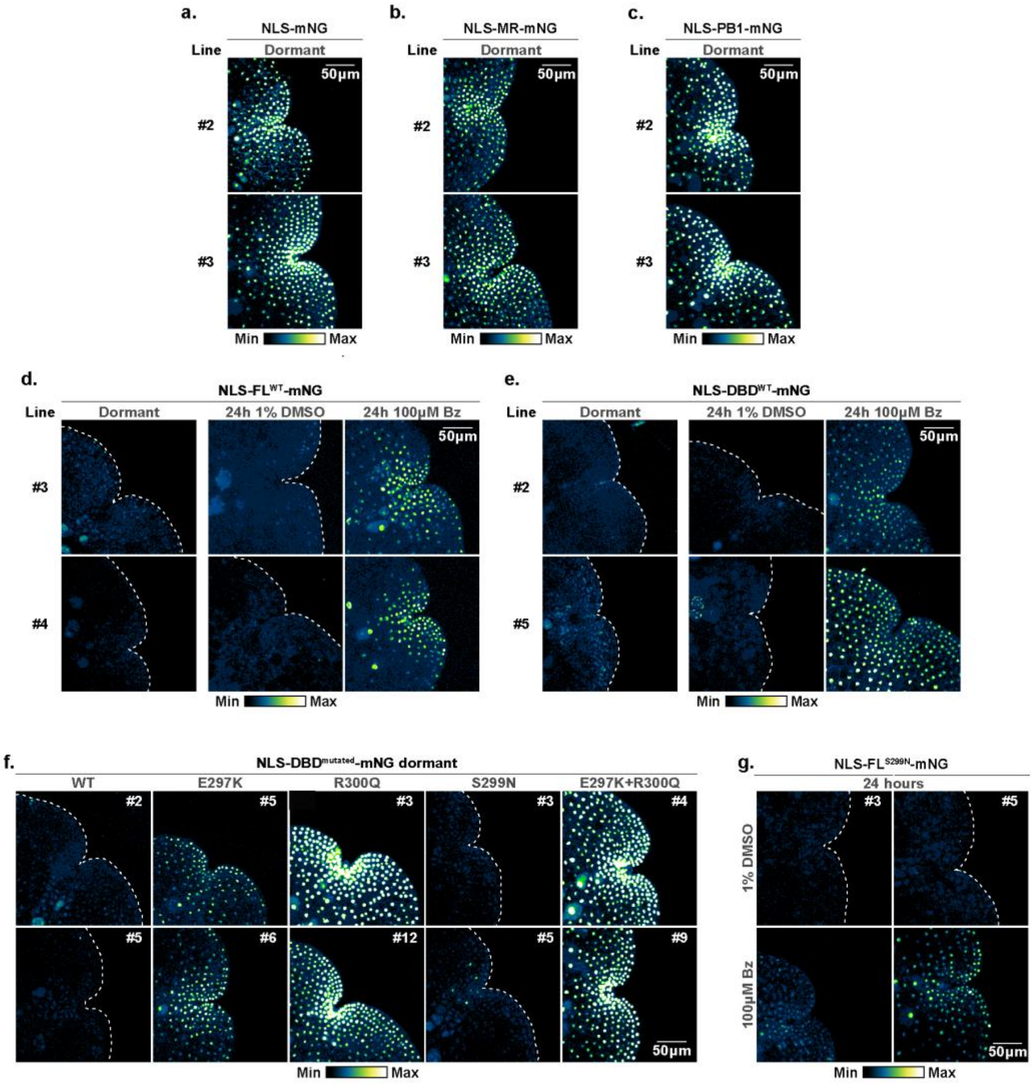


**Supplementary Figure 1: Assessment of stability fusion proteins through confocal microscopy.**

**a.-g.** Confocal images of *M. polymorpha* gemmae, showing representative expression of different fusion proteins as outlined in Fig. 1d. Numbers denoted with a hash indicate independent transgenic lines, here we show protein behaviour in two additional independent lines for all panels in Fig. 1. Abbreviation Bz stands for Bortezomib.

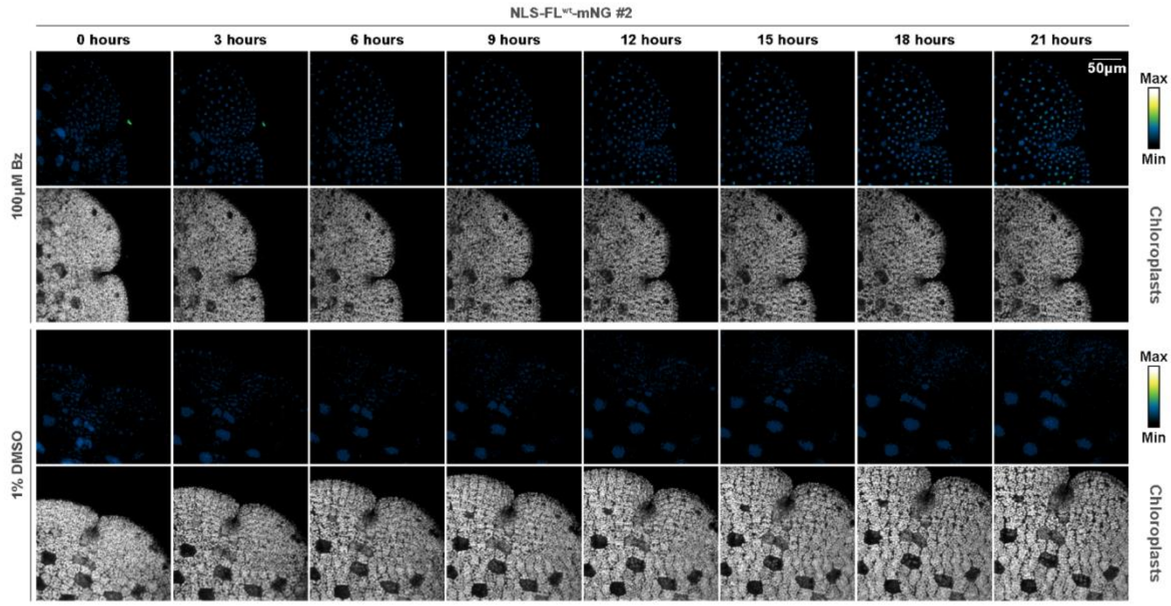


**Supplementary Figure 2: Dynamics and proteaseome-dependence of NLS-FL$^{wt}$-mNG accumulation**

Time series of gemma expressing NLS-FL$^{wt}$-mNG treated with Bortezomib (Bz) or DMSO for 21 hours.

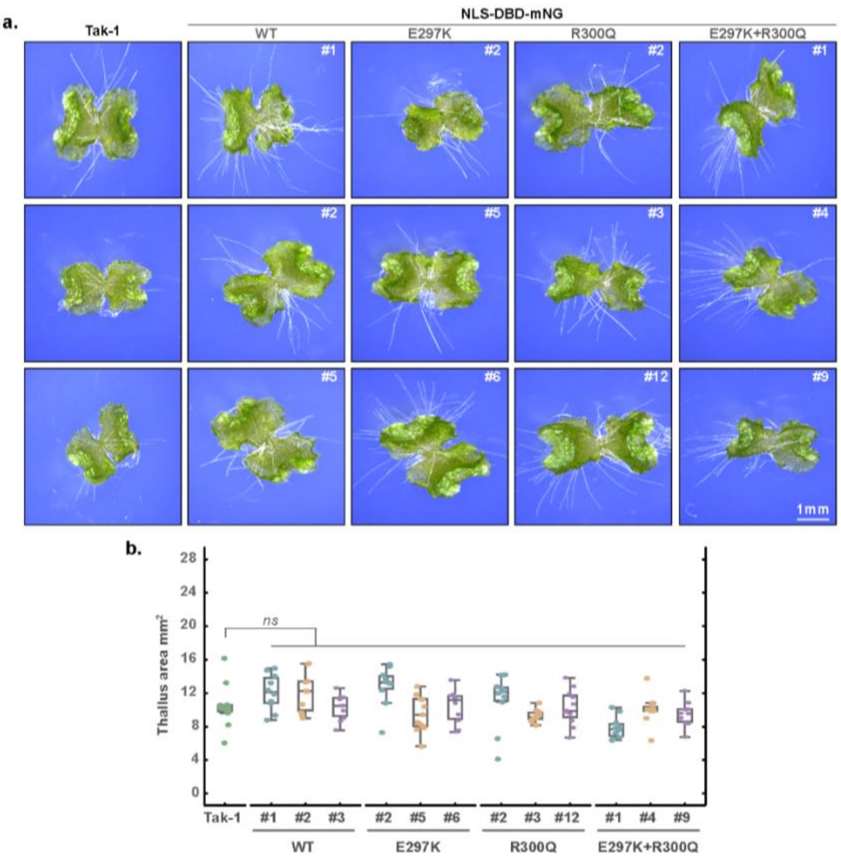


**Supplementary Figure 3: Expression of stabilized NLS-DBD-mNG fusions has no effect on Marchantia development**

**a.** Overview of phenotypes of seven day old gemmalings grown on ½ strength Gamborgs B5 medium wich express the NLS-DBD-mNG fusion proteins which harbour no mutations (WT) or mutations in key residues for stability as shown in Fig. 1h. **b.** Projected thallus area of aforementioned plants were measured and statisticaly compared by one-way ANOVA with Tukey post-hoc test (italics denote significant differences, $p$<0.05).

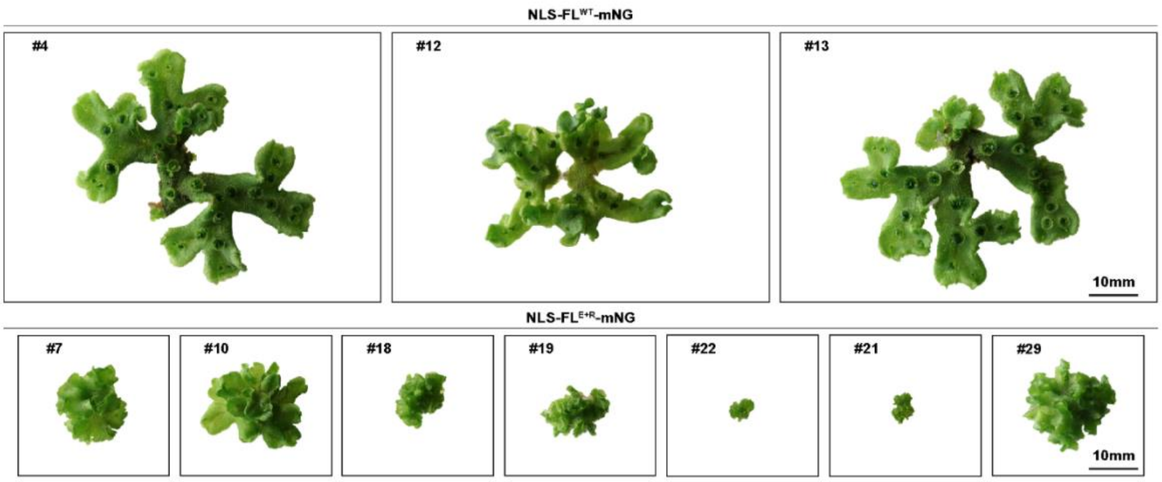


**Supplementary Figure 4: Expression of a stabilized full-length MpARF2 protein disturbs normal Marchantia development**

Overview of representative phenotypes of plants which survived $T_1$ selection on hygromycin-containing ½ Gamborgs B5 medium for 50 days. Compared here are multiple independent transgenic lines expressing NLS-FL$^{WT}$-mNG (top panel) and NLS-FL$^{E+R}$-mNG (bottom panel, harbours the E297K and R300Q mutations).

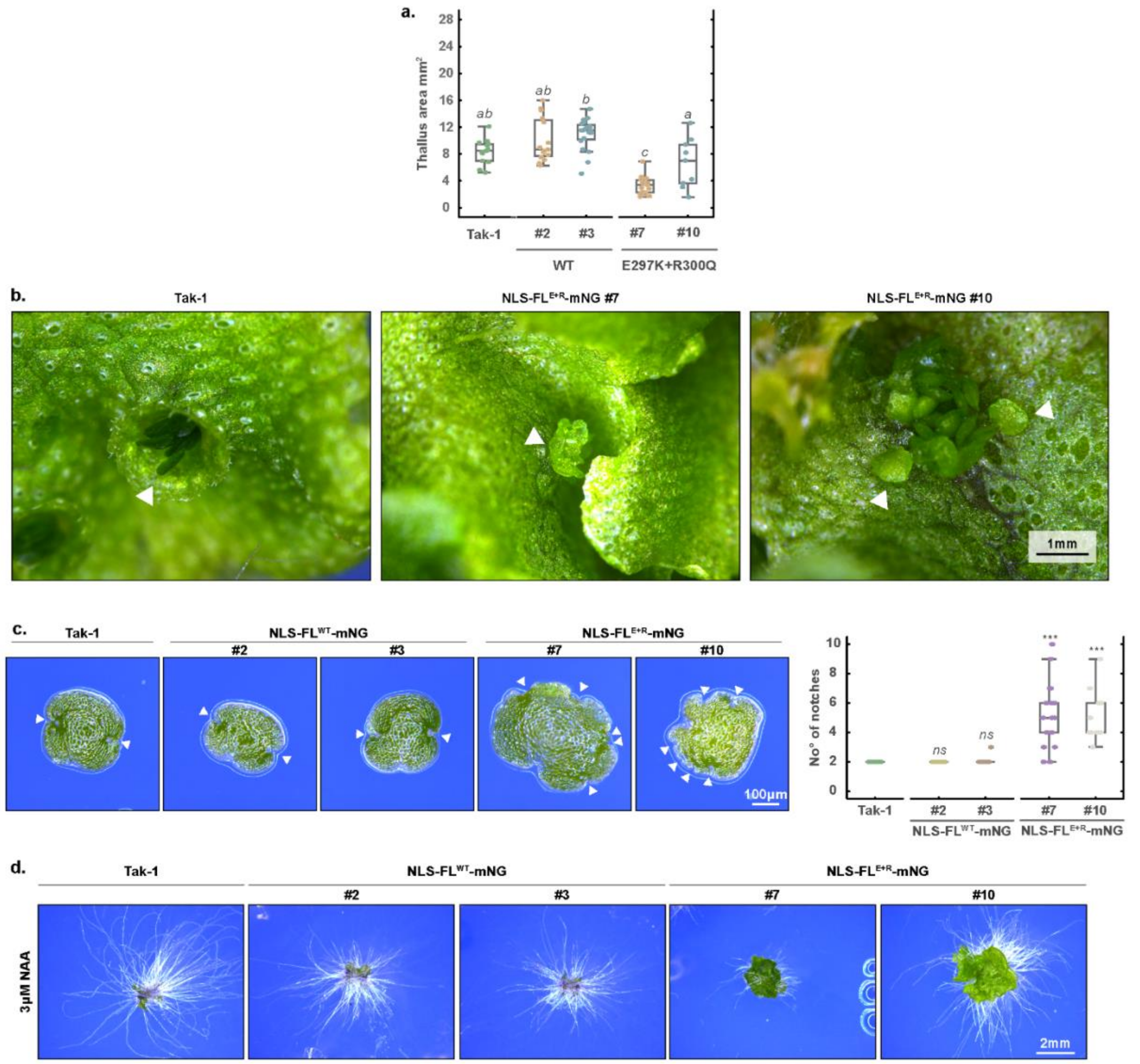


**Supplementary Figure 5: NLS-FL$^{E+R}$-mNG mutants exhibit pleitropic developmental defects and impaired auxin response**

**a.** Projected thallus area of seven day old gemmalings grown on ½ strength Gamborgs B5 medium. A comparison is made between the Tak-1 WT and plants wich express the NLS-FL$^{WT}$-mNG and NLS-FL$^{E+R}$-mNG fusion proteins (shown are two independent lines per construct). Thallus area was statisticaly compared by one-way ANOVA with Tukey post-hoc test (italics denote significant differences, $p<0.05$). **b.** Detailed images of gemmae formation in gemma cups (Tak-1, left) and on the thallus (NLS-FL$^{E+R}$-mNG #7 and #10, wherein hashes indicate independent transgenic lines). **c.** Stereomicroscope images of dormant gemmae of plants described in a. When clearly distinguishable apical notches were observed, these were counted and were statistically compared by a chi-squared test (*** $p<0.001$) to the Tak-1 WT. **d.** Overview of gemmalings of plants described in a. were grown on 3μM NAA containing ½ Gamborgs B5 medium for 17 days (same experiment as Fig. 2c but plants were grown for an additional seven days).

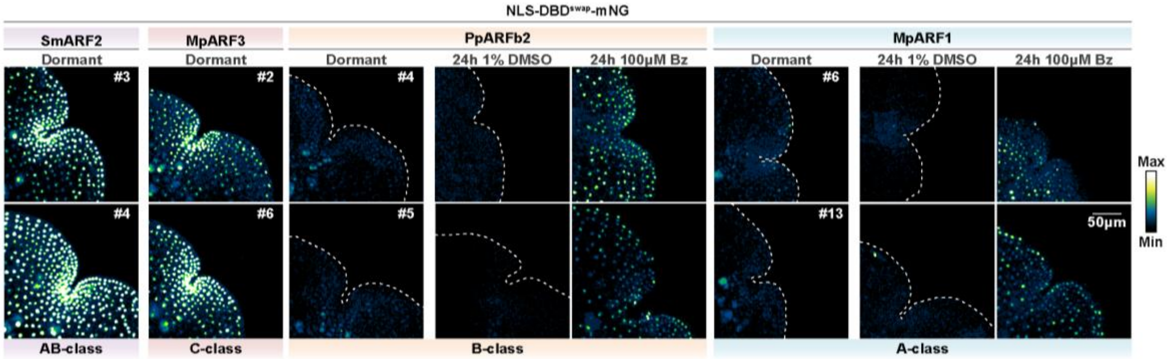


**Supplementary Figure 6: Investigation of the ability of homologous regions of ARF proteins from different streptophyte species in conferring instability.**

Confocal images of *M. polymorpha* gemmae, showing expression of different NLS-DBD-mNG fusion proteins with homologous sequences of other ARFs swapped. Numbers denoted with a hash indicate independent transgenic lines, here we show protein behaviour in two additional independent lines for all panels in Fig. 1. Bz stands for Bortezomib.

### Supplementary Table 1

| Primer | Sequence 5' > 3' | Description |
|---|---|---|
| **MdR298** | TGCATGCCTGCAGGTCGACTAAAAGCCTGTAATCACACGACG | Fw promoter *MpARF2* |
| **MdR299** | GCTGTCTAGAGGTCGGAACTTCTGTCTAAAATGC | Rv promoter *MpARF2* |
| **MdR306** | TTAGACAGAAGTTCCGACCTATGTCAGAAGCATCTTCCA | Fw A2 CDS to clone into pA2_XbaI_mNG Hifi |
| **MdR314** | ATGCTGCCGCCGCCAAGCTGCATGTCGTCGCCGCGCG | Rv PB1 to clone into pA2_XbaI_mNG |
| **MdR308** | CCATGCGGCAACGAAGAAATCTCGATT | Fw DBD to introduce E297K mutation and 2 fragment Hifi into XbaI site of pA2_XbaI_mNG together with a PB1 rev |
| **MdR309** | AATCGAGATTTCTTCGTTGCCGCATGG | Rv DBD to introduce E297K mutation and 2 fragment Hifi into XbaI site of pA2_XbaI_mNG together with a A2 start fw |
| **MdR310** | CGGAGAAATCTCAATTCTCTCTAATTT | Fw DBD to introduce R300Q mutation and 2 fragment Hifi into XbaI site of pA2_XbaI_mNG together with a PB1 rev |
| **MdR311** | AAATTAGAGAGAATTGAGATTTCTCCG | Rv DBD to introduce R300Q mutation and 2 fragment Hifi into XbaI site of pA2_XbaI_mNG together with a A2 start fw |
| **MdR312** | ATGCGGCAACGAAGAAATCTCAATTCTCTCT | Fw DBD to introduce E+R mutation (E297K + R269Q) and 2 fragment Hifi into XbaI site of pA2_XbaI_mNG together with a PB1 rev |
| **MdR313** | AGAGAGAATTGAGATTTCTTCGTTGCCGCAT | Rv DBD to introduce E+R mutation (E297K + R269Q) and 2 fragment Hifi into XbaI site of pA2_XbaI_mNG together with a A2 start fw |
| **MdR412** | GCGGCAACGGAGAAAAATCGATTCTCTCTAA | Fw introduce S299N in MpARF2 and 2 fragment Hifi into XbaI site of pA2_XbaI_mNG together with a PB1 rev |
| **MdR413** | TTAGAGAGAATCGATTTTTCTCCGTTGCCGC | Rv introduce S299N in MpARF2 and 2 fragment Hifi into XbaI site of pA2_XbaI_mNG together with a A2 start fw |
| **MdR315** | ATGCTGCCGCCGCCAAGCTGGAAGGGCTCCACTTCCCATG | Rv A2 AD (DBD end) to clone into pA2_XbaI_mNG |
| **MdR317** | ATGGCTCCAAAGAAGAAGAGAAAGGTC | Fw A2 DBD to add SV40 NLS at N-terminus |
| **MdR318** | TTAGACAGAAGTTCCGACCTATGGCTCCAAAGAAGAAG | Fw SV40 NLS to clone into XbaI of plasmid pGWB100 pA2_XbaI_mNG |
| **MdR319** | GGTTGTAAATTAGAGAGAATTGAGATTTCTCCGTTGCCGC | Rv Rv DBD to make R300Q mutation and 2 fragment Hifi into XbaI pA2_XbaI_mNG together with A2 start fw |
| **MdR320** | GCGGCAACGGAGAAATCTCAATTCTCTCTAATTTACAACCC | Fw Fw DBD to make R300Q mutation and 2 fragment Hifi into XbaI pA2_XbaI_mNG together with PB1 rev |
| **MdR372** | TTAGACAGAAGTTCCGACCTATGGCTCCAAAGAAGAAGAGAAAGG | Fw SV40 hifi overhang into xbaI with nls |
| **MdR373** | ATGCTGCCGCCGCCAAGCTGAACTGGACCTTGTTGGAAATTTTGCG ACG | Rv MpA2 MR hifi overhang into xbal with nls |
| **MdR374** | ATGCTGCCGCCGCCAAGCTGCATGTCGTCGCCGCGCGCCC | Rv MpA2 PB1 hifi overhang into xbal with nls |
| **MdR381** | TGGTTGTCCCTGCACTGCCAGATGGGCAGCGGCCGCGAGAAC | Rv swap degron MpARF3 into MpARF2 DBD longer overhang |
| **MdR382** | CCCATCTGGCAGTGCAGGGACAACCATTCTCTCTAATTTACAACCCT CG | Fw swap degron MpARF3 into MpARF2 DBD |
| **MdR389** | CCCATGCGGCAATTTCGGGATCTACATTCTCTCTAATTTACAACCCTC GATC | Fw swap degron SmARF2 into MpARF2 DBD longer overhang |
| **MdR390** | AATGTAGATCCCGAAATTGCCGCATGGGCAGCGGCCGCGAGAACC CC | Rv swap degron SmARF2 into MpARF2 DBD longer overhang |

| | | |
|---|---|---|
| **MdR353** | TTCCCTCCCTTGACGCATGGGCAGCGGCCGCGAGAA | Rv swap degron PpARF2 into MpARF2 DBD |
| **MdR354** | CCATGCGTCAAGGGAGGGAATGCGATTCTCTCTAATTTACAACCC | Fw swap degron PpARF2 into MpARF2 DBD |
| **MdR349** | GAATCGAGAATTCGTCGCTGCCGCATGGGCAGCGGCC | Rv swap degron MpARF1 into MpARF2 DBD |
| **MdR350** | CCGCTGCCCATGCGGCAGCGACGAATTCTCGATTCTCTCTAATTTACAACCC | Fw swap degron MpARF1 into MpARF2 DBD |
